# Sphingomyelinase disables Piezo1 channel inactivation to enable sustained response to mechanical force

**DOI:** 10.1101/792564

**Authors:** Jian Shi, Adam J Hyman, Dario De Vecchis, Jiehan Chong, Laeticia Lichtenstein, T. Simon Futers, Myriam Rouahi, Anne Negre Salvayre, Nathalie Auge, Antreas C Kalli, David J Beech

**Author notes:** To whom correspondence should be addressed: Professor David Beech, Leeds Institute of Cardiovascular and Metabolic Medicine, LIGHT Building, Clarendon Way, School of Medicine, University of Leeds, Leeds LS2 9JT, UK. Dr Jian Shi, Leeds Institute of Cardiovascular and Metabolic Medicine, LIGHT Building, Clarendon Way, School of Medicine, University of Leeds, Leeds LS2 9JT, UK. Dr Antreas Kalli, Leeds Institute of Cardiovascular and Metabolic Medicine, LIGHT Building, Clarendon Way, School of Medicine, University of Leeds, Leeds LS2 9JT, UK.

## Abstract

Piezo1 channels are determinants of vascular responses to fluid flow. They importantly provide sustained response to flow, but this property contrasts with the rapid inactivation that has become a hallmark of the channels in heterologous overexpression studies. Here we reveal a mechanism by which blood vessels disable inactivation to enable sustained physiological response. Creation of a molecular model of Piezo1 channel in defined lipid membranes suggested potential modulation by sphingomyelin and its product ceramide. Biological relevance was indicated by the observation that exogenous sphingomyelinase enhanced Piezo1-mediated Ca^2+^ entry in cultured endothelial cells. We therefore hypothesised that endogenous sphingomyelinase suppresses channel inactivation. Remarkably, in endothelium freshly-isolated from murine artery, neutral sphingomyelinase inhibitors or genetic disruption of sphingomyelin phosphodiesterase 3 (SMPD3) caused flow- and pressure-activated Piezo1 channels to become inactivating. SMPD3 retained its ability to disable inactivation in cell-free membrane patches, providing evidence for a membrane localised effect. The data suggest that inherent inactivation of Piezo1 channels is disabled by enzymatic control of lipid environment to enable physiological response to mechanical force.

## INTRODUCTION

Shear stress is a frictional force that arises when fluid flows along a cellular surface: it impacts many aspects of biology^1^. Especially rapid flow occurs in the vasculature where powerful shear stress arises from blood or lymph flowing against endothelium^2–4^. The ability of endothelial cells to sense this force and respond appropriately is critical in vascular development, health and disease^1,4–7^. Yet the underlying molecular mechanisms (particularly how the force is sensed) have been difficult to elucidate^4,7–10^. In 2014 an important molecular component was revealed as the Piezo1 channel^11–13^, a Ca^2+^ -permeable non-selective cationic channel that seems to have as its primary function the sensing of mechanical force and its transduction into cellular response^14–17^. Endothelial Piezo1 is relevant to vascular maturation in the embryo, response to shear stress in adult endothelial cells in vitro and in vivo, and the disease of generalized lymphatic dysplasia^11–13,18–25^. It alone confers shear stress response on otherwise resistant cells and responds quickly and reversibly in membrane patches excised from endothelial cells, suggesting a membrane-delimited, perhaps direct, ability to sense shear stress^11,12,18,19^. It does not inactivate in response to sustained flow and so impacts continuously until flow ceases^11,19,24,25^. Here we focus on the non-inactivating property because it contrasts with the rapid inactivation seen when Piezo1 is over-expressed in cell lines^14,17,26,27^.

## RESULTS

### Predicted interaction of Piezo1 with sphingomyelin and ceramide

Molecular simulation of Piezo1 channel was generated using published structural data^28^ and incorporated in model membranes of defined composition. The simulation predicted Piezo1 interaction with the membrane constituent, sphingomyelin (Figure 1a, b). Sphingomyelinase catalyses the generation of ceramide from sphingomyelin^29^. Therefore, we also analysed the channel in the presence of ceramide (5%) and a combination of ceramide and sphingomyelin (2.5% each); interactions with both lipids were indicated (Figure 1c, d). Some of the C-terminal amino acid residues predicted to interact with ceramide and sphingomyelin are close to determinants of inactivation^27^, suggesting the possibility to modulate inactivation (Figure 1e). We also analysed the dome-like membrane indentation caused by Piezo1 channel because it is thought to be a key property regulating channel gating^28,30^. Intriguingly, the depth of the dome depended on the relative proportions of ceramide and sphingomyelin (Supplementary Figure S1). In 5% sphingomyelin (0% ceramide) the dome depth was 5.62 ± 1.00 nm and 5.23 ± 1.08 nm for the upper and lower leaflets, whereas in 5% ceramide (0% sphingomyelin) the depth was 5.29 ± 0.79 nm and 4.94 ± 0.86 nm for the upper and lower leaflets. The analysis predicts importance of sphingomyelin and ceramide for Piezo1 and potential for regulation of inactivation by these lipids. We therefore used sphingomyelinase to alter the proportions of these lipids in native cells.

**Figure 1.**
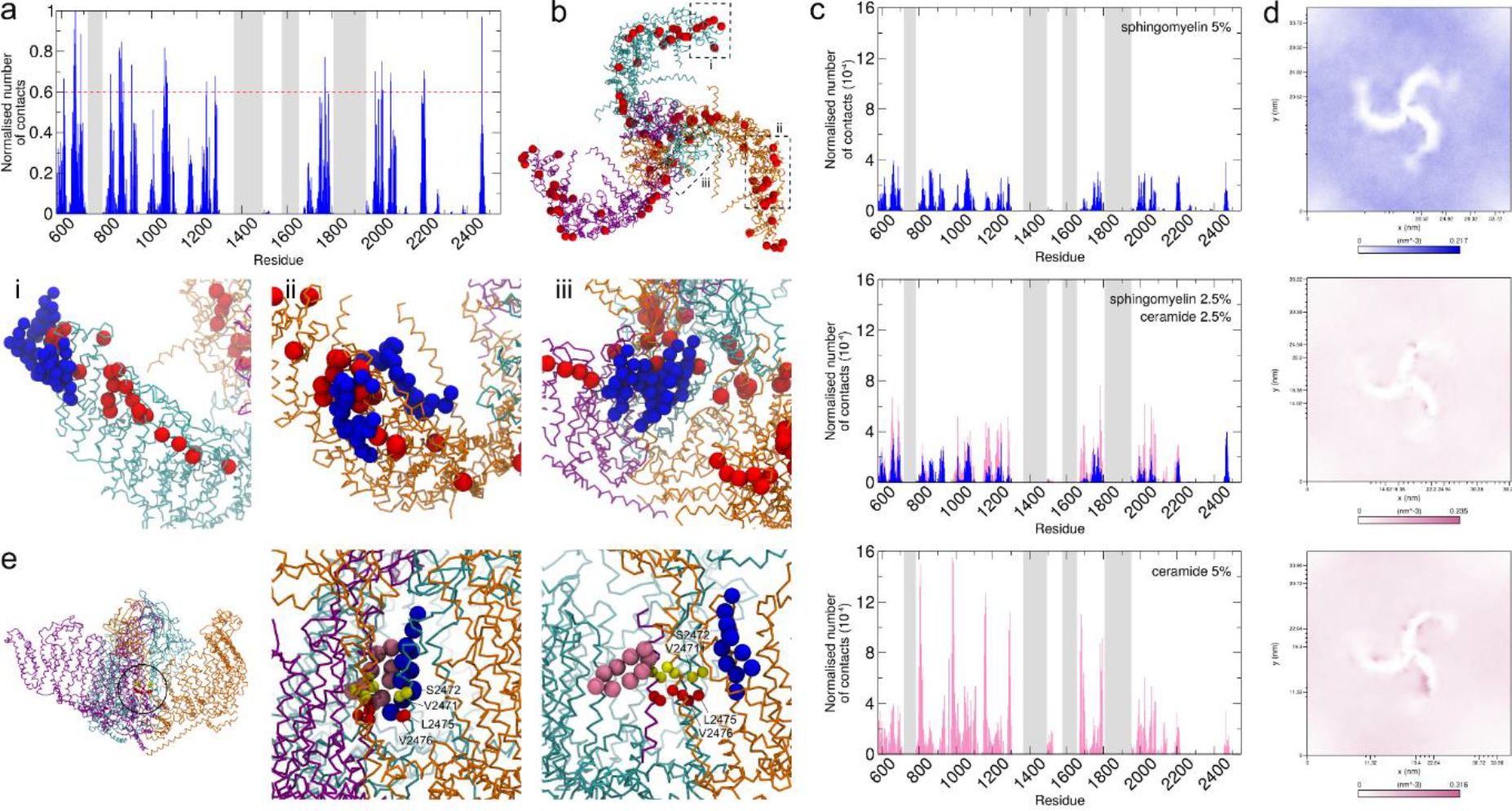
Predicted interaction of Piezo1 with sphingomyelin and ceramide. (**a**) Normalized average (over all repeat simulations) number of contacts (within a cut-off distance of 0.55 nm) between Piezo1 and the lipid sphingomyelin. The number of contacts of the 3 Piezo1 subunits (shown in orange, cyan and purple in (**b**) were added together. In the histogram, 0 represents no contacts and 1 the maximum number of contacts. The grey bars represent missing regions from the Piezo1 model. (**b**) Snapshot from one simulation performed with Piezo1 in a native bilayer (view from the top). The backbone particles of the residues above 0.6 in (**a**) are highlighted as red spheres. Detail of interactions are magnified in **i**, **ii** and **iii**. Sphingomyelin molecules that were in contact with the region that is magnified are shown in blue. (**c**) Normalized average number of contacts between Piezo1 and sphingomyelin (blue) and ceramide (pink) lipids in the bilayer. For the normalization, the number of contacts of each residue with a lipid type was divided by the number of lipids and the number of frames. (**d**) 2-dimensional average density plots for sphingomyelin (blue) and ceramide (pink). (**e**) Side view of Piezo1 structure. The circle indicates the locations of magnified snapshots on the right from two different simulations and shown in different orientations. Some of the amino acid residues predicted to interact with sphingomyelin and ceramide (yellow spheres: S2472, V2471) are close to suggested determinants of inactivation (red spheres: L2475, V2476). Sphingomyelin and ceramide molecules are shown as blue and pink spheres respectively.

### Bacterial sphingomyelinase enhances Piezo1-dependent Ca^2+^ entry

We first studied cultured human umbilical vein endothelial cells (HUVECs) as a model of native endothelium. These cells express functional Piezo1 channels^11,31^. A synthetic chemical agonist, Yoda1^31,32^, activated the channels to cause Ca^2+^ entry (Figure 2a). The response to Yoda1 was suppressed by Gd^3+^, a non-specific Piezo1 channel blocker^14^ or by depletion of Piezo1 caused by Piezo1-targeted siRNA, consistent with Piezo1 channels mediating the response (Figure 2a-c). To hydrolyse endogenous sphingomyelin, bacterial sphingomyelinase (bSMase) was added. It enhanced the Yoda1 response (Figure 2a-c). Heat-inactivated bSMase lacked effect (Figure 2d). Active bSMase lacked effect on other Ca^2+^ responses evoked by ATP or VEGF, suggesting specificity (Figure 2e) but there was a Piezo1-independent effect on baseline Ca^2+^ (Figure 2c). The data suggest ability of sphingomyelinase to positively regulate Piezo1 channels.

**Figure 2.**
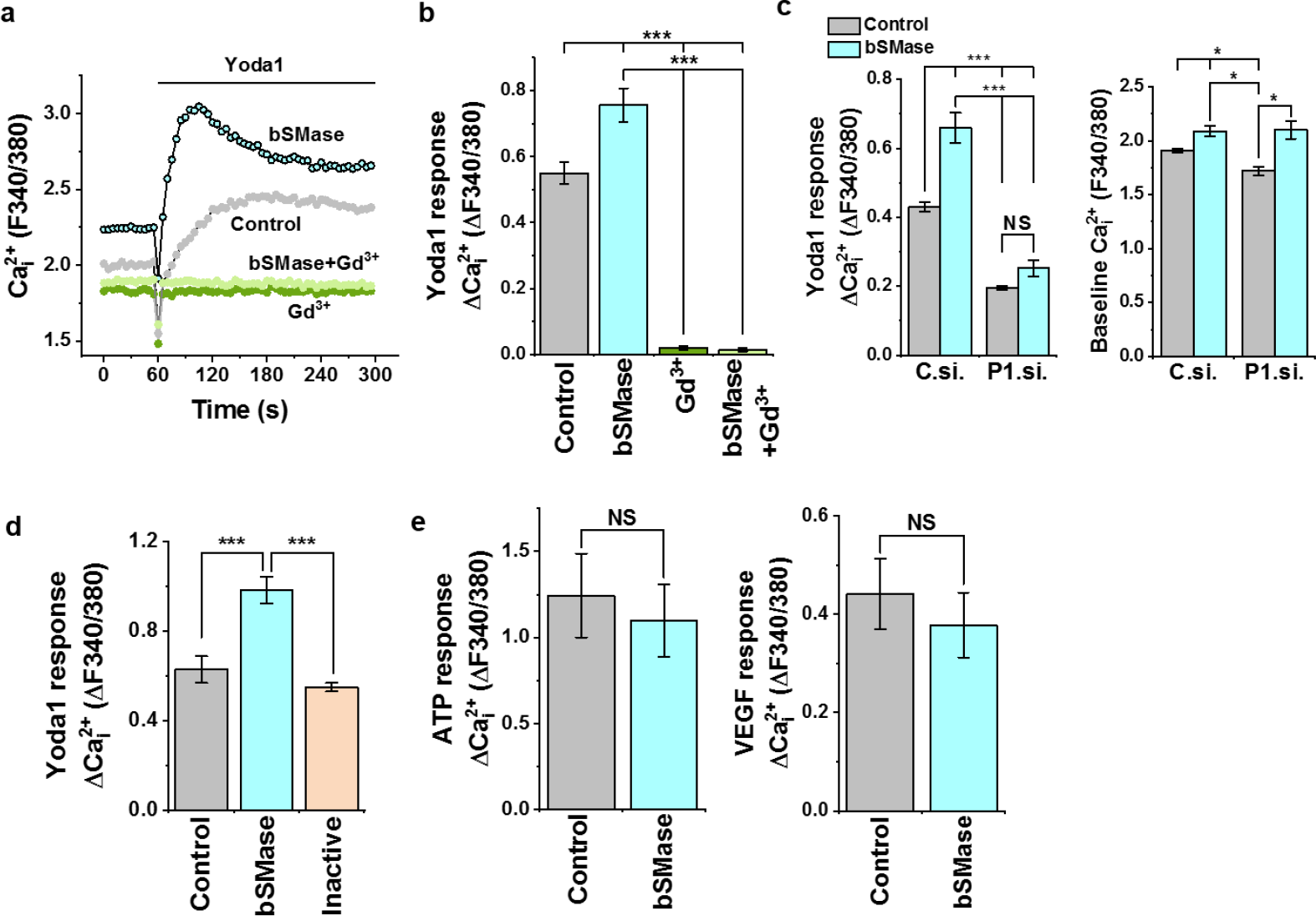
Bacterial sphingomyelinase enhances Piezo1 function. Data are intracellular Ca^2+^ measurements from human umbilical vein endothelial cells (HUVECs). (**a**) Example time-series traces showing effects of 1 μM Yoda1 on intracellular Ca^2+^ indicated by F340/380 in cells in a single 96-well plate without pre-treatment (Control) or after pre-treatment for 30 min at 37 °C with 0.5 U.mL^−1^ bacterial sphingomyelinase (bSMase) or 100 μM gadolinium chloride (Gd^3+^) or both. (**b**) As for (**a**) but plotted as the change in intracellular Ca^2+^ (ΔCa^2+^_i_) in response to Yoda1 and showing mean ± s.e.mean data for multiple independent experiments (n/N=3/4) where *** indicates *P* < 0.01. (**c**) Mean ± s.e.mean data for ΔCa^2+^_i_ in response to 1 μM Yoda1 (left) and baseline Ca^2+^ indicated by basal F340/380 in the same experiments (n/N=3/4) where *** indicates *P* < 0.01, * P < 0.05 and NS not significantly different. C.si. was control siRNA and P1.si. Piezo1 siRNA. (**d**) Similar to (**b**) but comparing the effect of 30 min pre-treatment with 0.5 U.mL^−1^ bSMase or the same concentration of heat-inactivated bSMase on the effect of 1 μM Yoda1 (n/N=3/4, *** *P* < 0.01). (**e**) Similar to (**b**) but in which Ca^2+^ responses were evoked by 20 μM ATP or 30 ng.mL^−1^ VEGF in place of Yoda1 (n/N=3/4 each, NS not significantly different).

### Neutral sphingomyelinase inhibitors prevent sustained endothelial response to flow

To investigate flow responses of native channels, endothelium was freshly isolated from second-order mesenteric arteries of adult mice. Our previous membrane potential recordings from this preparation showed an essential role for endogenous Piezo1 in determining resting membrane potential and sustained depolarization evoked by fluid flow^19^. Here we made similar observations, again observing robust reversible depolarization in response to fluid flow (Figure 3a). Small oscillations, of unknown mechanism, variably occurred superimposed on the overall response (Figure 3a). To test the contribution of endogenous sphingomyelinase we first incubated cells with pharmacological inhibitors of two different subtypes of this enzyme. Desipramine, a weak base that inhibits acid sphingomyelinase^33^, had no effect on the flow response, but two inhibitors of neutral sphingomyelinase (GW4869 and altenusin^29,34,35^) caused flow responses to become transient, such that sustained response was no longer evident (Figure 3b-f). There was sustained response only in the vehicle-control condition or the presence of desipramine (Figure 3e, f). None of the agents affected the resting membrane potential (Figure 3e). The data suggested that a neutral sphingomyelinase was required for the sustained response to flow.

**Figure 3.**
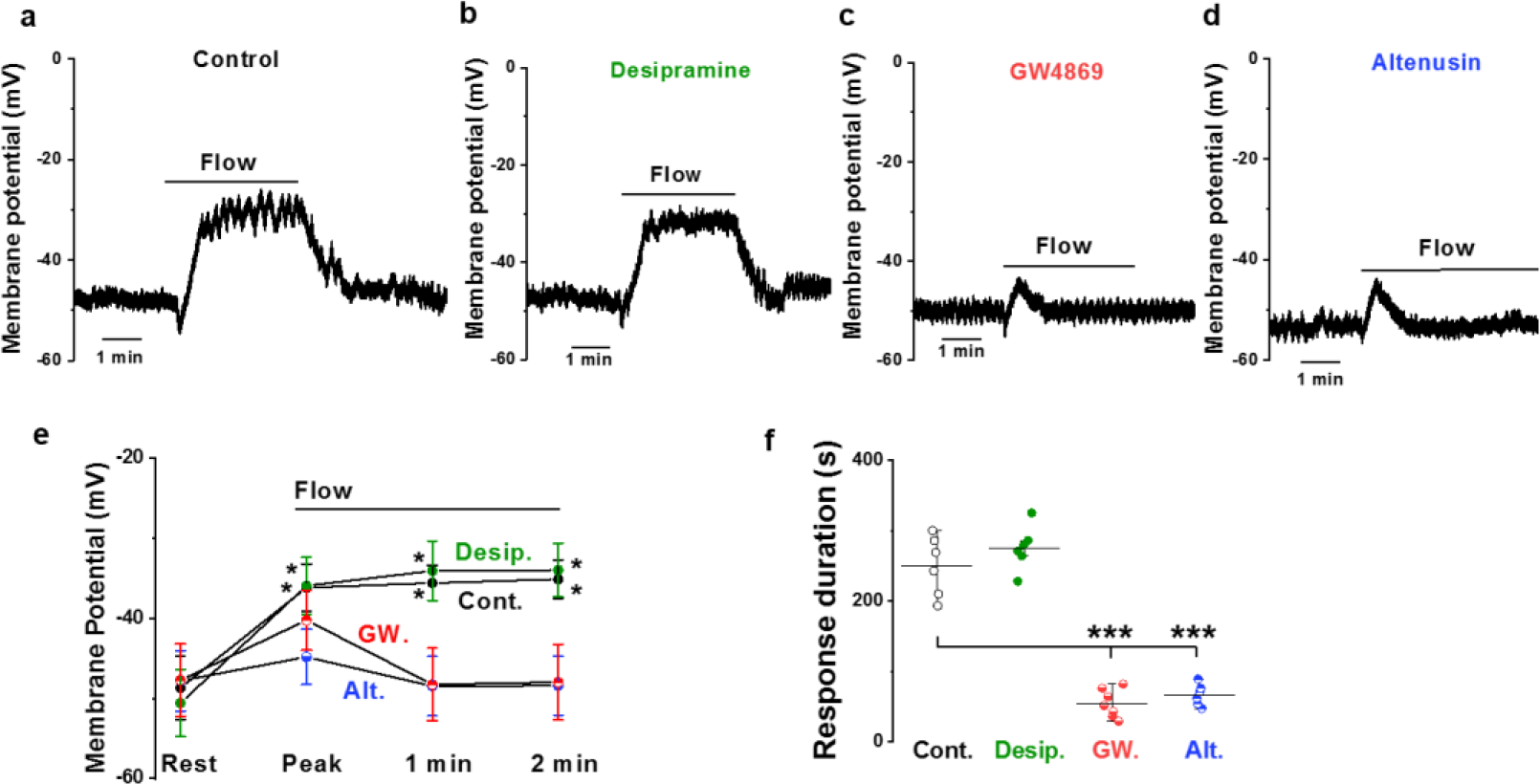
Inhibitors of neutral sphingomyelinase suppress sustained flow-evoked depolarisation. All data relate to membrane potential recorded in amphotericin whole-cell mode from freshly isolated endothelium of second-order mesenteric arteries. (**a**) Representative recording showing the sustained response to application of 20 μL.s^−1^ fluid flow in the presence of the vehicle control (2.5% DMSO). (**b-d**) Similar to (**a**) but showing responses after 10-min pre-incubation and the continuous presence of 10 μM Desipramine (**b**), 10 μM GW4869 (**c**) or 10 μM Altenusin (**d**). (**e**) Mean ± s.e.mean data for experiments of the type shown in (**a**-**d**): Control (Cont., n=6 recordings); Desipramine (Desip., n=6); GW4869 (GW., n=7); Altenusin (Alt., n=6). ‘Rest’ indicates the resting membrane potential in static (no flow) condition. ‘Peak’ indicates the maximum depolarisation of the membrane potential in response to flow (i.e. the initial response). Membrane potential values 1 and 2 min after the start of flow are also shown. (**f**) For the experiments of (**e**), the duration of the flow-evoked depolarisation. In Cont. and Desp., the duration of the response was the same as the duration of the exposure to flow. Mean ± s.e.mean data * *P* < 0.05, \*\*\**P* < 0.001.

### Disruption of *Smpd3* prevents sustained response to flow

Small-molecule inhibitors are not necessarily specific and do not shed light on the subtype of neutral sphingomyelinase, so we sought to identify the underlying gene. A prominent neutral sphingomyelinase in vascular biology is known to be sphingomyelin phosphodiesterase 3 (SMPD3 or nSMase2)^36^. It is encoded by the *Smpd3* gene, the best-characterised of the four nSMase genes^29,36^. We therefore investigated mice homozygous for the fragilitas ossium (Fro) mutation of the *Smpd3* gene (*Smpd3*^*fro*/*fro*^), which causes loss of the C-terminal active domain of SMPD3^29,37–39^. Strikingly, there was an initial transient response but no sustained response to flow in these mice (Figure 4a-d). The data suggest that SMPD3 is required for sustained flow-evoked depolarization. The resting potential was not significantly changed by *Smpd3*^*fro*/*fro*^ (Figure 4c). The data suggest that SMPD3 is required for the sustained response to flow.

**Figure 4.**
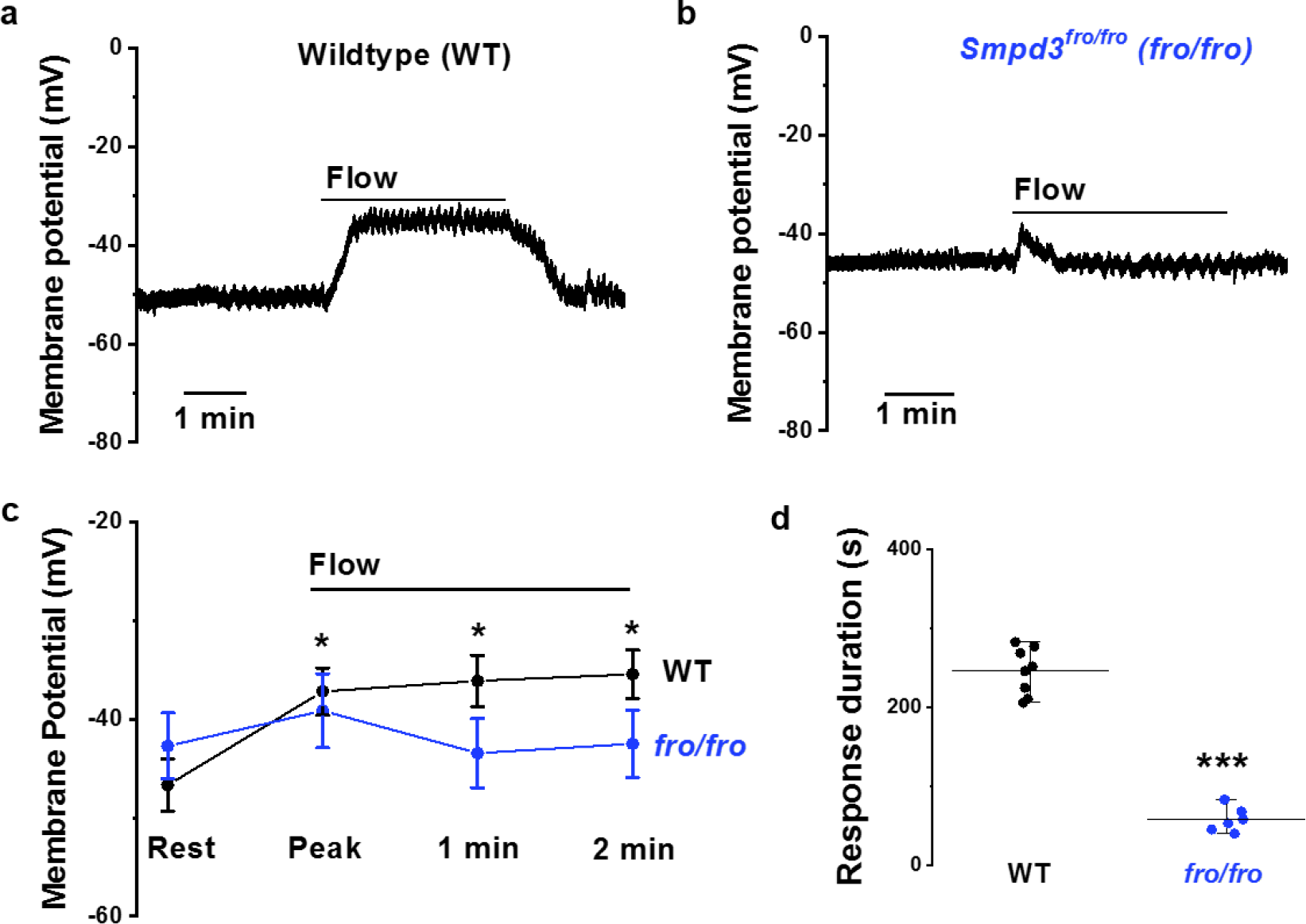
Absence of sustained flow-evoked depolarisation in *Smpd3*^*fro*/*fro*^ mice. All data relate to membrane potential recorded in amphotericin whole-cell mode from freshly isolated endothelium of second-order mesenteric arteries. (**a**) Representative recording showing response to application of 20 μL.s^−1^ fluid flow in endothelium of wildtype (WT) mice. (**b**) Representative recording showing response to application of 20 μL.s^−1^ fluid flow in endothelium of *Smpd3*^*fro*/*fro*^ mice (*fro/fro*). (**c**) Mean ± s.e.mean data for experiments of the type shown in (**a**, **b**). ‘Rest’ indicates the resting membrane potential in static (no flow) condition. ‘Peak’ indicates the maximum depolarisation of the membrane potential in response to flow, which was the initial response. Membrane potential values 1 and 2 min after the start of flow are also shown. (**d**) For the experiments of (**c**), the duration of the flow-evoked depolarisation. In the WT group, the duration of the response was the same as the duration of the exposure to flow. Mean ± s.e.mean data * *P* < 0.05, \*\*\**P* < 0.001. WT: n=8; *Fro*/*fro*: n=6 recordings.

### Regulation by SMPD3 is local and membrane-delimited

Prior work suggests that sustained activation of Piezo1 channels is preserved in membrane patches excised from endothelium, suggesting that intracellular communication and organelles are not required^19^. All channel activity evoked by fluid flow in these studies was Piezo1 dependent and had the expected unitary current properties of Piezo1 channels^19^. It is also known that SMPD3 is a plasma membrane-tethered enzyme^29^. Therefore we investigated if the SMPD3-Piezo1 relationship is functional in excised cell-free membrane. Similar to the membrane potential observations from intact endothelium (Figure 4), *Smpd3*^*fro*/*fro*^ caused Piezo1 channels to become inactivating (Figure 5a-c). Consistent with these data, the observed channel activity had the expected unitary current size of Piezo1 channels at −80 mV^15,18,19^ and was suppressed by Gd^3+^, a blocker of Piezo1 channels^14,19^ (Figure 5d-f). GW4869 similarly suppressed the sustained response to flow in excised patches (Supplementary Figure S2). The data suggest that SMPD3 operates locally to suppress inactivation of Piezo1 channels.

**Figure 5.**
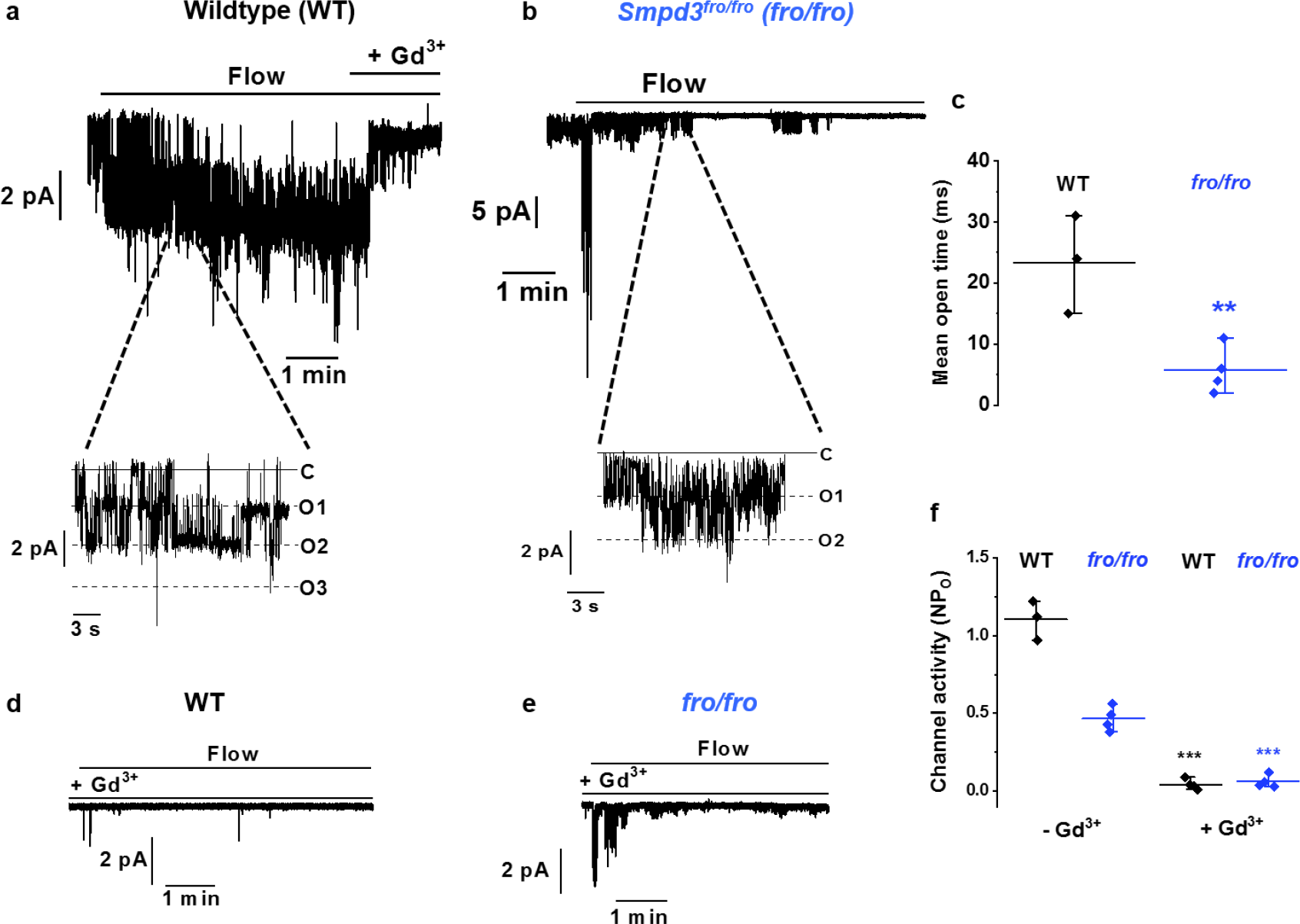
Piezo1 and SMPD3 function together in a membrane-delimited mechanism. All measurements were single channel current recordings from outside-out patches excised from freshly isolated endothelium of second-order mesenteric arteries. Holding potential was −80 mV. (**a-b**) Representative recordings showing responses to 20 μL.s^−1^ fluid flow in endothelium from wildtype (WT) mice (**a**) or *Smpd3*^*fro*/*fro*^ (*fro*/*fro*) mice (**b**). Note the large initial peak current in the upper panel of (**b**). (**c**) For experiments of the type shown in (**a**-**b**), all data points and mean ± s.e.mean for the channel open time 20 s after the initiation of flow (i.e. the sustained response) (WT n=3, *fro*/*fro* n=4). ** *P* < 0.01. (**d**-**e**) As for (**a**-**b**) but in the presence of 10 μM Gd^3+^. (**f**) For experiments of the type shown in (**a**, **b**, **d**, **e**), all data points and mean ± s.e.mean for channel activity represented by NP_o_ (channel number multiplied by open probability) measured 20 s after the initiation of flow. Without Gd^3+^ (- Gd^3+^): WT n=3, *fro*/*fro* n=4 recordings. With Gd^3+^ (+ Gd^3+^): WT n=4, *fro*/*fro* n=4 recordings. *** *P* < 0.001.

### SMPD3 prevents pressure-evoked inactivation

Activation by flow is physiological but slower in onset than the rapid force caused by pressure pulses and cell-prodding used in studies of overexpressed Piezo1 channels^14,26,27^. We therefore also applied pressure pulses to cell-attached patches of endothelium. As previously described in HUVECs^11^ there was sustained channel activation in response to 20-mmHg pressure pulses applied to freshly isolated endothelium (Figure 6a). The currents were large multi-channel currents, larger than those seen in outside-out patches (Figure 6a *cf* Figure 5a) perhaps at least partly because larger patch pipettes were used for these recordings. Strikingly, in cell-attached patches on endothelium of *Smpd3*^*fro*/*fro*^ mice the current inactivated, as shown by strong decay of the inward current despite the persistent pressure step (Figure 6b). Unexpectedly the initial peak current was larger than the control (Figure 6c) but it then decayed to become much smaller than the control (Figure 6d). GW4869 similarly caused the current to inactivate strongly but did not cause an increase in amplitude (Supplementary Figure S3), suggesting that the larger current in *Smpd3*^*fro*/*fro*^ mice reflects compensatory increase in channel density. The data suggest that SMPD3 also prevents inactivation when the mechanical stimulus is a pressure pulse.

**Figure 6.**
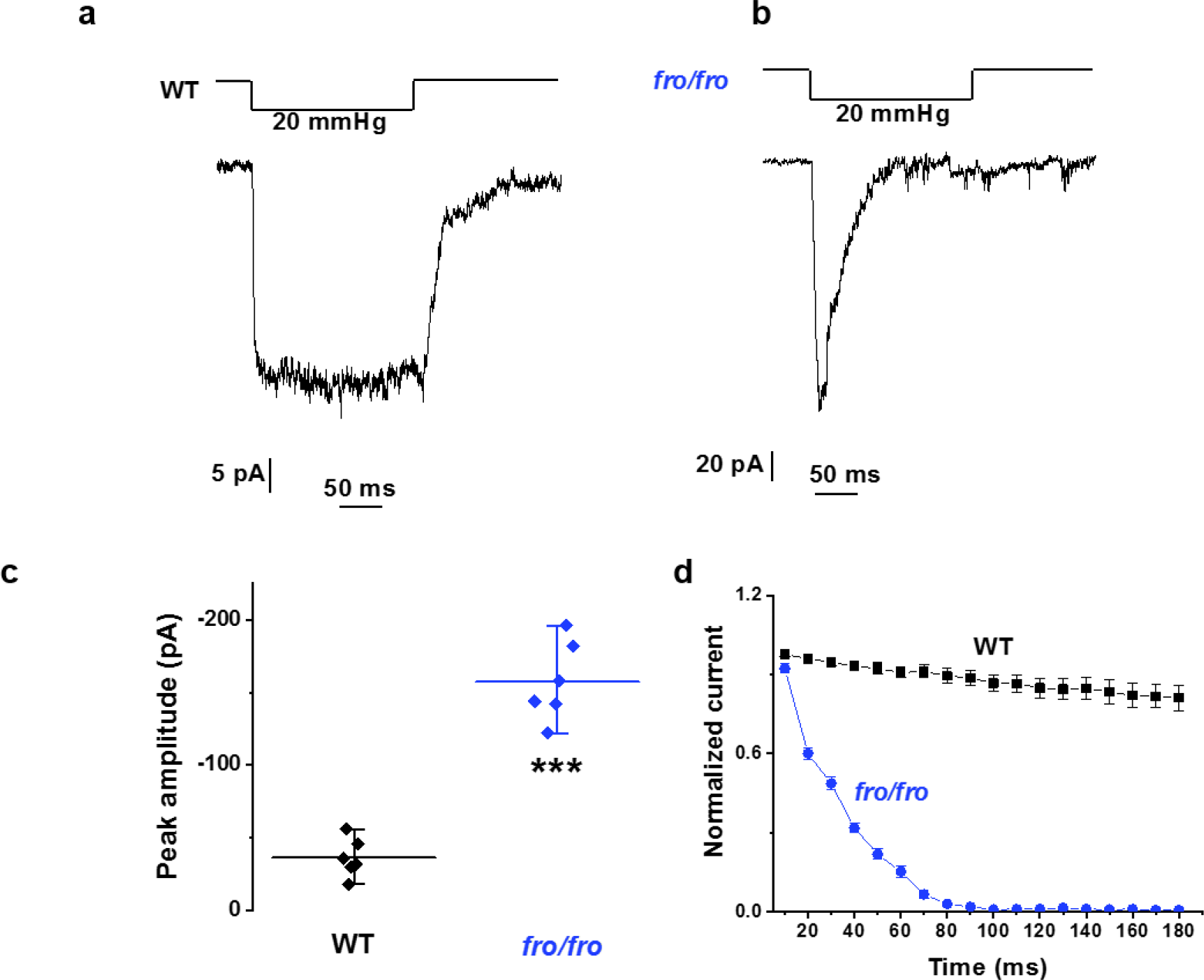
SMPD3 is also important when the stimulus is a pressure pulse. All measurements were made from cell-attached patches on freshly isolated endothelium of second-order mesenteric arteries of adult mice. (**a-b**) Representative recordings showing responses to 20 mmHg negative pressure steps of 200 ms duration for endothelium from wildtype (WT) mice (**a**) and *Smpd3*^*fro*/*fro*^ (*fro*/*fro*) mice (**b**). (**c**) Mean ± s.e.mean data for the peak (maximum) inward current amplitude that occurred in experiments of the type shown in (**a**, **b**): WT recordings (n=6 recordings); *Smpd3*^*fro*/*fro*^ (*fro*/*fro*) recordings (n=6). *** *P* < 0.001. (**d**) As for (c) but showing analysis of the current decay rate for WT (n=6) and *Smpd3*^*fro*/*fro*^ (*fro*/*fro*) (n=6) recordings.

## DISCUSSION

Based on this study we suggest that endothelial cells use an enzymatic mechanism to alter the local lipid environment of Piezo1 channels to prevent inherent inactivation and allow physiologically important mechanical response. Our findings are consistent with a study pre-dating the discovery of Piezo1 that suggested a role for SMPD3 in endothelial response to fluid flow^40^. Lipid regulation of inactivation is also a reasonable proposal because the inactivation gate is hydrophobic^27^ and sphingomyelin and ceramide are predicted to achieve proximity to this gate (Figure 1e). Moreover, prior work has shown that application of lipids, such as fatty acids, to Piezo1 channels suppresses their inactivation^41^. We envisage that SMPD3 achieves its effect through constitutive activity that determines the local lipid profile but its activity may also be enhanced by Ca^2+^ entry because SMPD3 locates to the inner leaflet of the bilayer and interacts with calcineurin, a Ca^2+^ - and calmodulin-dependent serine/threonine protein phosphatase that activates SMPD3 by regulating its phosphorylation status^29,42^.

Surprisingly, only mechanically-activated channel activity was regulated by SMPD3, not constitutive activity. This suggests two pools of channel: one that avoids inactivation through SMPD3 and the other that avoids it through a different mechanism (as yet unidentified). We speculate that the constitutively active channel exists in a lipid raft that confers a compartment for constitutive neutralisation of inactivation, so that these channels are unaffected by SMPD3 disruption because they are already non-inactivating.

It will be worthwhile to investigate if SMPD3-dependent escape from inactivation is also important in non-endothelial cell types where rapid inactivation is also likely to be incompatible with function. One possibility is osteoblasts where non-inactivating Piezo1 activity has been seen^43^. It may not be a coincidence that *Piezo1*- and *Smpd3*-disrupted mice both show abnormal bone formation^38,43^. Slow inactivation has also been described in other cell types such as embryonic stem cells, where a role of Piezo1 in the relatively slow process of cell proliferation has been suggested^44^.

It has been reported previously that addition of exogenous sphingomyelinase modifies gating properties of voltage-gated ion channels, for example slowing inactivation of voltage-gated Na^+^ channels depending on host cell type^45,46^. These findings are consistent with our observations but, to the best of our knowledge, ours is the first indication that an endogenous sphingomyelinase regulates an ion channel for context-specific modulation. It will be worthwhile to investigate the broader relevance of this concept because of the known importance of sphingomyelin and ceramide in membranes, the widespread expression of sphingomyelinases and the diversity of ion channels in complex membranes.

## METHODS

### Piezo1 channel modelling

Structural data were obtained from the mouse cryo-EM structure PDB: 6B3R^28^. Missing residues were added with MODELLER (v 9.19)^47,48^ and the loop refinement tool was used to remove a knot in one chain between residues 1490-1498. The best loop was selected out of 10 candidates according to the discrete optimized protein energy method^49^. The final Piezo1 mouse model does not comprises the first N-terminal 576 residues and residues 718-781, 1366-1492, 1579-1654, 1808-1951. Therefore, each chain is composed by five non-overlapping fragments: residues 577-717, 782-1365, 1493-1578, 1655-1807 and 1952-2547. The Piezo1 model obtained was further energy minimized in vacuum with GROMACS 5.0.7^50^ prior simulations.

### Coarse-grained simulations

The Piezo1 model obtained as described above was converted to a coarse-grained (CG) resolution and energy minimised. The CG molecular dynamics (CG-MD) simulations were performed using the Martini 2.2 force field^51,52^ and GROMACS 5.0.7^50^. In the Martini force field, there is an approximate 4:1 mapping of heavy atoms to coarse-grained particles. To model the protein secondary and tertiary structure an elastic network model with a cut-off distance of 7 Å was used. The elastic network restricts any major conformational change within the protein during the CG-MD simulations. Further, the model was inserted in a complex asymmetric bilayer using the INSert membrANE tool^53^. For the upper leaflet a concentration of 55% 1-palmitoyl-2-oleyl-phosphtidylcholine (POPC), 5% sphingomyelin, 20% 1-palmitoyl-2-oleyl-phosphtidylethanolamine (POPE), and 20% cholesterol, was used. For the lower leaflet a concentration of 50% POPC, 20% POPE, 5% sphingomyelin, 20% cholesterol, 5% 1-palmitoyl-2-oleyl-phosphtidylserine, and 5% phosphatidylinositol 4,5-bisphosphate, was used. Further simulations were performed substituting half of the sphingomyelin in the upper leaflet (i.e. 2.5%) with ceramide or by depleting all sphingomyelin (i.e. 5%) with ceramide. The systems were neutralized with 150 mM NaCl. The model was further energy minimized and subsequently equilibrated for 500 ns with the protein particles restrained (1000 kJ.mol^−1^.nm^−2^) to allow the membrane bilayer to equilibrate around the protein. After each equilibration, five unrestrained repeat simulations of 5 μs each were run for every system starting from different velocities. Both equilibration and production runs were performed at 323 K which is above the transition temperature of all the lipid species used. Protein, lipids and solvent were separately coupled to an external bath using the V-rescale thermostat^54^ (coupling constant of 1.0). Pressure was semi-isotropically maintained at 1 bar (coupling constant of 1.0) with compressibility of 3 × 10-6 using the Berendsen^55^ and the Parrinello-Rahman^56^ barostats, for the equilibration and productions, respectively. Lennard-Jones and Coulombic interactions were shifted to zero between 9 and 12 Å, and between 0 and 12 Å, respectively.

### Trajectory analyses and molecular graphics

Contact analyses within the distance of 0.55 nm and lipid density around the protein were performed using the gmx mindist and gmx densmap tools from the GROMACS package. For the contacts analysis, all the beads of the lipid were considered. For the densities, all the five repeat simulations were concatenated and then fitted on the protein backbone particles. Molecular graphics were generated with the VMD 1.9.3^57^ (http://www.ks.uiuc.edu/Research/vmd/) and data were plotted using Grace (http://plasma-gate.weiz-mann.ac.il/Grace/).

### Quantification of dome depth

To obtain the dome depth, the simulation trajectory was fitted to the protein coordinates (reference structure). The coordinates of the CG phosphate particles in each frame of the fitted trajectory are extracted by a Python script. The phosphate particles are then used to separate the bilayer leaflets using a branching network algorithm. Briefly, this method starts with a single phosphate particle, and identifies the other phosphate particles which are within a cut-off distance (2 nm) of the starting particle. The cut-off distance is selected to be smaller than the separation between the bilayer leaflets. Phosphate particles identified in this way are grouped into the same leaflet as the starting residue. This process iterates repeatedly until no more new particles can be added, and the remaining particles are assumed to be part of the other leaflet. The process is then repeated starting in the other leaflet to validate the initial identification. For each leaflet, the depth of the dome is calculated as the difference between the surface level and the bottom of the dome. The surface level is taken to be the average z-coordinate for the phosphate particles with z-coordinate in excess of the 90th centile. The average is used here to minimise the effect of random fluctuation of the membrane. For the bottom of the dome, the z-coordinate of the phosphate particle with the absolute lowest z-coordinate is used. This is because the bottom of the dome is prone to far less fluctuation, being fixed to Piezo1, which in turn is the fitting reference for the rest of the simulation. To calculate the 2-dimentional height map of the z-coordinate of the CG phosphate particles in our simulations, for each frame CG phosphate particles from each leaflet were binned along the x and y axes; 75 bins for each axis. For each frame, the average z-coordinate of beads contained in each bin was calculated and stored in a matrix. The matrices of all frames are averaged to create the final height map, which is plotted using the PyPlot library. The code used is documented at: https://github.com/jiehanchong/membrane-depth-analysis.

### Wild-type and genetically modified mice

All animal use was authorized by the University of Leeds Animal Ethics Committee and The Home Office, UK. All animals were maintained in GM500 individually ventilated cages (Animal Care Systems) at 21°C 50-70% humidity, light/dark cycle 12/12 hr on RM1 diet (SpecialDiet Services, Witham, UK) ad libitum and bedding of Pure‘o Cell (Datesand, Manchester, UK). Genotypes were determined using real-time PCR with specific probes designed for each gene (Transnetyx, Cordova, TN). Male wild-type or *Smpd3*^*fro*/*fro*^ mice^39^ on 129/Sv background were 12-13 weeks old at the time of experiments. Otherwise, all mice were C57BL/6 males aged 10-14 weeks.

### Isolation of endothelium from mesenteric artery

Endothelium was freshly isolated from second-order branches of mouse mesenteric arteries as described previously^19^. Briefly, dissected second-order mesenteric arteries were enzymatically digested in dissociation solution (126 mM NaCl, 6 mM KCl, 10 mM Glucose, 11 mM HEPES, 1.2 mM MgCl_2_, 0.05 mM CaCl_2_, with pH titrated to 7.2) containing 1 mg.mL^−1^ collagenase Type IA (Sigma-Aldrich, Dorset, UK) for 14 min at 37 °C and then triturated gently to release endothelium on a glass coverslips for recordings on the same day.

### Patch-clamp electrophysiology

Recordings were made at room temperature using an Axopatch-200B amplifier equipped with a Digidata 1550A and pCLAMP 10.6 software (Molecular Devices, Sunnyvale, CA, USA). Endothelium was in a standard bath solution containing (mM) 135 NaCl, 4 KCl, 2 CaCl_2_, 1 MgCl_2_, 10 glucose and 10 HEPES (titrated to pH 7.4 using NaOH). For membrane potential recordings in zero current mode, heat-polished patch pipettes with tip resistances between 3 and 5 MΩ were used and contained amphotericin B (Sigma-Aldrich) as the perforating agent, added to a pipette solution containing (mM) 145 KCl, 1 MgCl_2_, 0.5 EGTA and 10 HEPES (titrated to pH 7.2 using KOH). Outside-out and cell-attached membrane patch recordings were made using the same equipment but in voltage-clamp mode. The tip resistances of recording pipettes for cell-attached recordings were between 3 and 5 MΩ, while for outside-out recordings they were between 12 and 15 MΩ. Currents were sampled at 20 kHz and filtered at 2 kHz. For cell-attached recordings, the extracellular (bath) solution contained (mM) 140 KCl, 1 MgCl_2_, 10 glucose and 10 HEPES (titrated to pH 7.3 using KOH), while the patch pipette contained (mM) 130 NaCl, 5 KCl, 1 CaCl_2_, 1 MgCl_2_, 10 Tetraethylammonium.Cl and 10 HEPES (titrated to pH 7.3 using NaOH). For outside-out recordings, the external solution was standard bath solution and the patch pipette contained (mM) 145 KCl, 1 MgCl_2_, 0.5 EGTA and 10 HEPES (titrated to pH 7.2 using KOH). For application of fluid flow, endothelium or a membrane patch was manoeuvred to the exit of a capillary tube with tip diameter of 350 μm, out of which ionic (bath) solution flowed at 20 μL.s^−1^. For pressure pulses, 0.2-s duration square pulses were applied to the patch pipette every 5 s using a High Speed Pressure Clamp HSPC-1 System (ALA Scientific Instruments, USA).

### Culture and transfection of human umbilical vein endothelial cells (HUVECs)

HUVECs were purchased from Lonza and cultured in Endothelial Cell Basal Medium (EBM-2) supplemented with 2% foetal calf serum and: 10 ng.mL^−1^ vascular endothelial growth factor (VEGF), 5 ng.mL^−1^ human basic fibroblast growth factor, 1 μg.mL^−1^ hydrocortisone, 50 ng.mL^−1^ gentamicin, 50 ng.mL^−1^ amphotericin B and 10 μg.ml^−1^ heparin (BulletKitTM, Lonza). Experiments were performed on cells from passage 2-5. HUVECs were transfected with 20 nM siRNA using Lipofectamine 2000 in OptiMEM (Gibco) as per the manufacturer’s instructions (Invitrogen). Medium was replaced after 4-5 hr and cells were used for experimentation 48 hr post-transfection. Piezo1 siRNA was GCCUCGUGGUCUACAAGAUtt (Ambion), which was previously validated^11^. Non-targeting control siRNA was from Dharmacon.

### Intracellular Ca^2+^ measurement

Measurements were made at room temperature on a FlexStation 3 (Molecular Devices, California) bench-top fluorometer controlled by Softmax Pro software v5.4.5 (Figure 2.1). Cells were plated in clear 96-well plates (Corning, NY, USA) at a confluence of 90% 24 hours before experimentation. Cells were incubated with fura-2AM (2 μM) (Molecular ProbesTM) in SBS containing 0.01% pluronic acid (Thermo Fisher Scientific) for 60 min at 37 °C. Cells were washed with SBS for 30 min at room temperature. Baseline fluorescence ratios were recorded before the addition of the compound solution to the cell plate after 60 seconds, with recording thereafter for a total of 300 seconds. The Standard Bath Solution (SBS) contained: NaCl 135 mM, KCl 5 mM, MgCl_2_ 1.2 mM, CaCl_2_ 1.5 mM, D-glucose 8 mM and HEPES 10 mM. pH was titrated to 7.4 using 4M NaOH.

### Chemicals and reagents

Unless stated otherwise, all commercially available chemicals and enzymes were purchased from Sigma-Aldrich (Dorset, UK). Yoda1 and desipramine was purchased from Tocris (Bio-Techne Ltd., Abingdon, UK).

### Data analysis

Genotypes of mice were always blinded to the experimenter and mice were studied in random order determined by the genotype of litters. Data were generated in pairs (control mice and *Smpd3*^*fro*/*fro*^ mice) and data sets compared statistically by independent t-test without assuming equal variance. Paired t-tests were used when comparing data before and after application of flow or a substance to the same membrane patch or cell. One-way ANOVA followed by Tukey posthoc test was used for comparing multiple groups. Statistical significance was considered to exist at probability (P) <0.05 (*<0.05, **<0.01, ***<0.001). Where data comparisons lack an asterisk, they were not significantly different. The number of independent experiments (mice or independent cell cultures) is indicated by n. For multi-well assays or multiple cell on coverslip studies, the number of replicates is indicated by N. Descriptive statistics are shown as mean±s.e.mean. Origin Pro software was used for data analysis and presentation.

## DATA AVAILABILITY

All relevant data are available from the authors upon reasonable request.

## ACKNOWLEDGEMENTS

The research was supported by a British Heart Foundation Intermediate Research Fellowship to JS, Wellcome Trust Investigator and British Heart Foundation programme grants to DJB, an Academy of Medical Sciences and Wellcome Trust Springboard Award to AK and INSERM funding to MR, ANS and NA. AJH was supported by a BBSRC PhD Studentship and JC by a MRC Clinical Research Training fellowship. We thank Katsuhiko Muraki and Mark Kearney for helpful feedback on the manuscript.

## AUTHOR CONTRIBUTIONS

JS performed and analysed patch-clamp recordings and isolated endothelial cells. AJH cultured endothelial cells and performed intracellular Ca^2+^ measurements. DDV and ACK generated the Piezo1 molecular model. DDV performed and analysed the simulations. JC wrote the script for analysing dome depth. LL, TSF, MR, ANS and NA prepared genetically-modified mice. DJB, JS and ACK raised funds for the project, generated ideas, interpreted data and co-wrote the paper. DJB and JS organised the project.

## ADDITIONAL INFORMATION

Supplementary molecular modelling and data figures accompany this paper (Supplementary Figures S1, S2 and S3).

