## Supplemental Data for "Sphingomyelinase disables Piezo1 channel inactivation to enable sustained response to mechanical force"

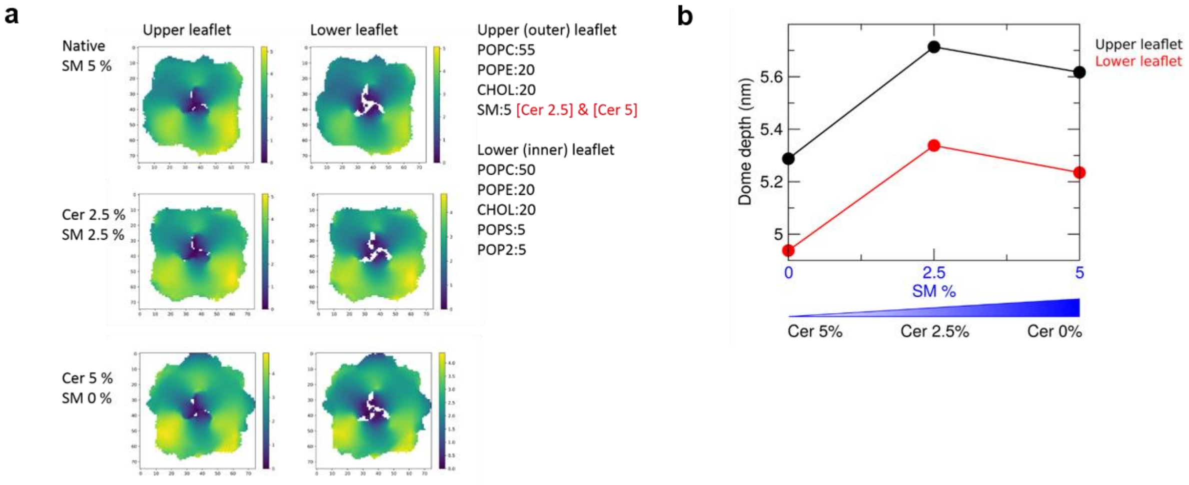


**Supplementary Figure S1. Predicted impact of sphingomyelin and ceramide on the Piezo1 channel dome.** (**a**) Height map of the z-coordinate of the CG phosphate particles corresponding to the upper (left) and lower (right) leaflet, averaged across all repeat simulations of the same simulation system. This analysis is shown for the 3 different simulation systems with different concentrations of sphingomyelin and ceramide. (**b**) Average dome depth of the upper and lower leaflets in the 3 simulation systems. With 2.5% sphingomyelin, the dome reached maximum depth of 5.71 ± 0.87 nm and 5.34 ± 0.95 nm for the upper and lower leaflets.

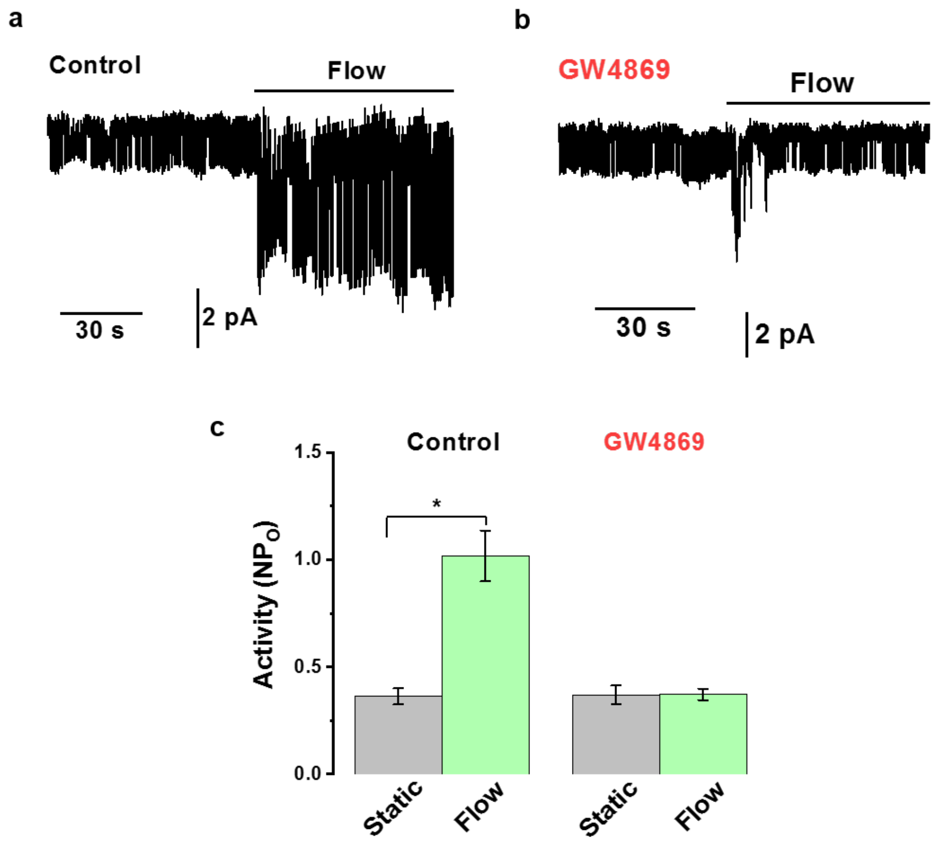


**Supplementary Figure S2. GW4869 prevents sustained channel activation by flow.** (**a**, **b**) Example traces for Piezo1 channel currents in outside-out patches voltage-clamped at -80 mV. Constitutive channel activity was observed prior to application of flow at 20 μL.s$^{-1}$, which stimulated multiple channel openings. (**a**) Control condition: vehicle (2.5% DMSO). (**b**) Test condition: 10 μM GW4869. (**c**) Mean ± s.e.mean channel activity data for experiments of the type shown in (**a**, **b**): Control (n=4 recordings); GW4869 (n=4). NP$_o$ is the number of channels multiplied by the open probability of the channels. * $P < 0.05$ by Student's t-test.

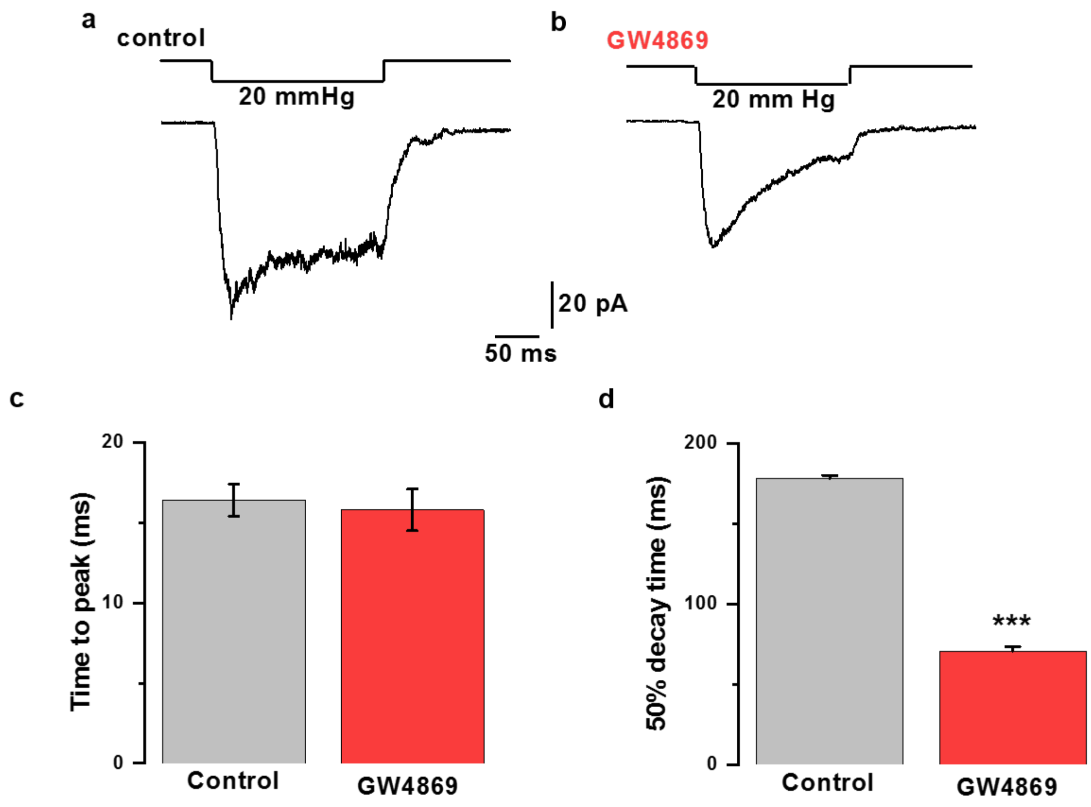


**Supplementary Figure S3. GW4869 prevents sustained channel activation by pressure step.** (**a**, **b**) Example traces for macroscopic Piezo1 channel currents in cell-attached patches voltage-clamped at -80 mV. Negative pressure steps of 20 mmHg were applied to the patch pipette. (**a**) Control condition: vehicle (2.5% DMSO). (**b**) Test condition: 10 µM GW4869. (**c**) Mean ± s.e.mean data for experiments of the type shown in (**a**, **b**), showing the time to reach peak inward current after first applying the pressure step: Control (n=6 recordings); GW4869 (n=6). (**d**) Mean ± s.e.mean data for experiments of the type shown in (**a**, **b**), showing the time for 50% decay of the current after reaching the peak: Control (n=6); GW4869 (n=6). *** $P < 0.001$ by Student's t-test,
